# Regulatory B cells (Bregs) inhibit osteoclastogenesis and prevent bone loss in osteoporotic mice model

**DOI:** 10.1101/2021.03.10.434751

**Authors:** Leena Sapra, Asha Bhardwaj, Pradyumna K. Mishra, Bhupendra K. Verma, Rupesh K. Srivastava

## Abstract

Increasing evidences in recent years have suggested that regulatory B cells (Bregs) are crucial modulator in various inflammatory disease conditions. However, the role of Bregs in case of postmenopausal osteoporosis remains unknown. Also, no study till date have ever investigated the significance of Bregs in modulating osteoclastogenesis. In the present study, we for the first time examined the anti-osteoclastogenic potential of Bregs under *in vitro* conditions and we observed that Bregs suppressed RANKL mediated osteoclastogenesis in bone marrow cells in a dose dependent manner. We further elucidated the mechanism behind the suppression of osteoclasts differentiation by Bregs and found that Bregs inhibit osteoclastogenesis via IL-10 production. To further confirm the bone health modulating potential of Bregs we employed post-menopausal osteoporotic mice model. Remarkably, our *in vivo* data clearly suggest a significant reduction (p < 0.01) in both CD19^+^IL-10^+^ and CD19^+^CD1d^hi^CD5^+^IL-10^+^ B10 Bregs in case of osteoporotic mice model. Moreover, our serum cytokine data further confirms the significant reduction of IL-10 levels in osteoporotic mice. Taken together, the present study for the first time unravels and establish the unexplored role of regulatory B cells in case of osteoporosis and provide new insight into Bregs biology in the context of post-menopausal osteoporosis.

## Introduction

B cells are classically characterized by their potential to produce and secrete antibodies. They also function as an antigen presenting cells (APCs) and produce several immunomodulatory cytokines. Nevertheless, B cells with immunosuppressive functions has also been observed in various conditions and these B cells are called as Regulatory B cells or “Bregs”. Microenvironment plays a crucial role in directing the development and differentiation of Bregs (Dar et al., 2020; Moufida et al., 2021). It has been observed that antigen and B cell receptor (BCR) signalling is crucial for the development of Bregs. In the presence of toll like receptor (TLR) ligands such as lipopolysaccharide (LPS), CpG and CD40 ligand the development of Bregs can be optimized (Tedder et al., 2015; Rosser et al., 2015). These B cells exhibits the ability to regulate various disease conditions including inflammatory bone loss diseases such as rheumatoid arthritis (RA), collagen induced arthritis (CIA) etc. (Yang et al., 2012; Flores-Borja et al., 2013). Bregs via production of interleukin (IL)-10, IL-35 and transforming growth factor (TGF)-β suppress the immunopathology by barring the expansion of various immune cells including T lymphocytes (Ran et al., 2020). Given that IL-10 producing B cells plays a crucial role in immune regulation, Tedder et al., defined a subset of Bregs named as “B10 cells” whose anti-inflammatory potential is only attributable to the production of IL-10 cytokine in various disease models such as cancer, autoimmune diseases and infectious diseases. The co-expression of CD1d and CD5 has later been utilized to characterize splenic B10 cells to inhibit the progression of inflammation upon stimulation for 5 hours with lipopolysaccharide (LPS), phorbal 12-myristate 13-acetate (PMA), ionomycin and monensin under *ex vivo* conditions (Yanaba et al., 2008; Yanaba et al., 2009). Various studies demonstrated that adoptive transfer of splenic B10 cells dampened the autoimmune reactions in several models of experimental autoimmune encephalitis (EAE), CIA, intestinal inflammation and systemic lupus erythematosus (SLE) (Matsushita et al., 2008; Watanabe et al., 2010; Yanaba et al., 2011; Yang et al., 2012). Thus, multiple studies in both humans and mice highlighted that Bregs suppress inflammatory reactions via IL-10 cytokine.

IL-10 is not just a signature cytokine for Bregs but also a potent modulator of immune response that also contributes towards the maintenance of bone health via inhibition of osteoclast mediated bone resorption (Qian et al., 2014). Various studies proposed that CD4^+^Foxp3^+^ Treg cells via secretion of IL-10 cytokine suppress osteoclastogenesis and thus bone resorption (Zaiss et al., 2007; Dar et al., 2018a, Dar et al., 2018b; Sapra et al., 2021). Interestingly, various studies reported that Breg cells mediates its immunosuppressive functions by inducing differentiation of naïve T cells into Tregs (Mielle et al., 2018). Nevertheless, the role of Bregs and its secretory cytokines on differentiation of osteoclasts has not been investigated till date. It has been also observed that numerical defect in the Tregs population along with its efficacy to produce IL-10 cytokine results in the development of various inflammatory bone loss conditions including osteoporosis. Osteoporosis is a systemic skeletal disease that is mainly characterized by the enhancement in the deterioration of bone tissue and low bone mass. Various factors act synergistically to enhance the likelihood of developing osteoporosis viz. age, sex, hormonal imbalance, dietary factors and immune system. Accumulating evidences lends support to the theory that bone destruction observed in osteoporosis caused by osteoclasts is due to impairment or loss of homeostatic balance of inflammatory and anti-inflammatory cells viz. Th17 and Tregs (Dar et al., 2018a, Dar et al, 2018b). With the growing involvement of immune system in the pathology of osteoporosis, our group has recently coined the term “Immunoporosis” (Srivastava et al., 2018). Among the global population, greater prevalence of osteoporosis has been observed in post-menopausal women due to deficiency of estrogen hormone. Surprisingly, it has also been found that estrogen hormone (E2) plays a crucial role in the development of Bregs during EAE (Benedek et al., 2016). E2 implantation showed neuroprotective effect in the brain of EAE mice by increasing the frequency of regulatory B cells. But the impact of estrogen deficiency on Bregs has not been thoroughly examined in case of post-menopausal osteoporosis.

In the present study we for the first-time report that under *in vitro* conditions CD19^+^IL-10^+^ and CD19^+^CD1d^hi^CD5^+^IL-10^+^ (B10) Bregs inhibit the differentiation of osteoclasts. In addition to this, we also for the first-time report that the frequencies of CD19^+^IL-10^+^ and B10 Bregs are significantly reduced in both bone marrow and spleen of ovariectomized induced post-menopausal osteoporotic mice model. Taken together our results further support that numerical defects in B10 Bregs along with its inability to secrete IL-10 cytokine *in vivo* may contribute towards the establishment of pro-inflammatory conditions in case of postmenopausal osteoporosis. Also, our results further attest and establish the important role of immune system in osteoporosis i.e., Immunoporosis.

## Materials and Methods

### Reagents and Antibodies

The following antibodies/kits were obtained from eBiosciences (USA): PerCp-Cy5.5 Anti-Mouse-CD19 (1D3) (45-0193-82), PE-Cy7 Anti-Mouse-CD5 (53-7.3) (25-0051-81), APC Anti-Mouse (CD1d) (1B1) (17-0011-82), Foxp3/Transcription factor staining buffer (0-5523-00) and RBC lysis buffer (00-4300-54). FITC Anti-Mouse–IL-10 (JES5-16E3) (505005) was purchased from Biolegend (USA). Protein transport inhibitor cocktail and Mouse TNF-α (560478) ELISA kit were procured from BD (USA). The following ELISA kits were brought from R&D: Mouse IL-10 (M1000B) and Mouse IL-17 (M1700). PMA, Ionomycin, LPS (*Escherichia coli* serotype 0111: B4) and Acid phosphatase leukocyte (TRAP) kit (387A) were procured from Sigma-Aldrich (USA). Macrophage-colony stimulating factor (M-CSF) (300-25) and Receptor activator of nuclear factor κB-ligand (sRANKL) (310-01) were procured from peprotech (USA). α-Minimal essential media (MEM) and RPMI-1640 were obtained from Gibco (Thermo Fisher Scientific, USA).

### Splenic B cell purification

Splenic B cells from C57BL/6 mice were purified by magnetic separation according to manufacturer’s instructions. Briefly, after RBC lysis, resulting cells were subjected to biotinylated mouse B cell enrichment cocktail (BD, USA) and incubated for 20-30 min at 4°C. Labelled cells were then washed carefully with washing buffer and incubated with streptavidin particles plus-DM for 30 min at 4°. Further, cells underwent magnetic separation using cell separation magnet and negative fraction comprised of resulting B cells were assessed for purity (>95 %) by flow cytometric analysis. Lastly, cells were suspended in RPMI-1640 media for *in vitro* cultures.

### B cells activation with LPS

Purified B cells were cultured in 24 well plate (2 × 10^6^) in 1ml/well in the presence and absence of LPS (10 ug/ml) for different time periods (5 hr and 24 hr) at 37°C in 5 % humidified CO2 incubator. At the end of incubation, activated B cells were harvested and washed thrice with 1XPBS containing 2 % FBS, especially to remove the traces of LPS before co-culturing with bone marrow cells for osteoclasts differentiation.

### Co-culture of LPS activated B cells with BMCs for Osteoclastogenesis

Generation of osteoclasts from mouse bone marrow cells (BMCs) was performed as previously described (Sapra et al., 2021). Briefly, BMCs were harvested from femur and tibiae of 8–10 wks old female mice and RBC lysis was performed with 1X RBC lysis buffer. After RBC lysis, cells were cultured overnight in T25 flask in endotoxin free α-MEM media supplemented with 10% heat inactivated fetal bovine serum (FBS) and M-CSF (35 ng/ml). On the following day, non-adherent cells (BMCs) (1×10^6^ cells/well) were collected and co-cultured with LPS activated and non-activated B cells in 24 well plate in osteoclastogenic medium supplemented with MCSF (30 ng/ml) and RANKL (100 ng/ml) for 4 days. At interval of 2 days old media was replenished with fresh media supplemented with fresh factors. After 4 days of incubation TRAP staining was carried out.

### Tartrate resistant acid phosphatase (TRAP) staining

For evaluating the generation of mature multinucleated osteoclasts tartrate resistant acid phosphatase (TRAP) staining was performed according to manufacturer’s instructions. Briefly, at the end of incubation cells were washed thrice with 1XPBS and fixed with fixative solution comprised of citrate, acetone and 3.7% formaldehyde for 10 min at 37 °C. After washing twice with 1X PBS, fixed cells were stained for TRAP at 37 °C in dark for 5–15 min. Multinucleated TRAP positive cells with≥3 nuclei were considered as mature osteoclasts. TRAP positive multinucleated cells were further counted and imaged using inverted microscope (ECLIPSE, TS100, Nikon). Area of TRAP positive cells was quantified with the help of Image J software (NIH, USA).

### Transwell experiment

Transwell chambers with 8 μM pore size membranes were employed to physically separate LPS-B cells from BMCs. Activated B cells were seeded in upper chamber and BMCs in the lower chamber at different cells ratio. After four days of incubation, cells were processed for evaluating osteoclastogenesis via TRAP staining.

### Flow Cytometric analysis of surface and intracellular markers

Cells were harvested and stained with antibodies specific for Bregs. For analysis of intracellular IL-10 cytokine by B cells, isolated splenocytes or purified B cells were resuspended in RPMI-1640 complete media comprised of 10% FBS and 0.1% mercaptoethanol (ME) at a density of 2 X10^6^ cells/ml in 24 well plate. Cells were then activated with LPS (10 ug/ml), PMA (50 ng/ml, Sigma Aldrich), Ionomycin (500 ng/ml, Sigma Aldrich) and protein transport inhibitor cocktail (BD, USA) for 5h. For surface marker staining, cells were first incubated with anti-CD19-PerCPcy5.5, anti-CD5-PE Cy7 and anti-CD1d-APC and incubated for 30 min in dark on ice. After washing, cells were fixed and permeabilized with 1X fixation-permeabilization-buffer for 30 min on ice in dark. Further, cells were stained with anti-IL-10-FITC for 45 min. After that cells were harvested and stained for IL-10 producing B cells. After washing, cells were acquired on BD FACS-Aria (USA). Flowjo-10 (TreeStar, USA) software was used to analyse the samples and gating strategy was done as per previously defined-protocols.

### Post-menopausal osteoporotic mice model

All *in vivo* experiments were carried out in 8-10 wks old female C57BL/6 mice. All the mice were housed under specific pathogen free conditions at the animal facility of All India Institute of Medical Sciences (AIIMS), New-Delhi-India. Mice were exposed to bilateral-ovariectomy after anesthetizing them with ketamine (100-150 mg/kg) and xylazine (5-16 mg/kg) intraperitoneally. Following groups were taken for the present study viz. Sham and Ovx (n=6/grp). Healthy control group (Sham) was subjected to sham surgery. Both the groups were maintained on a 12-h light/dark cycle in polycarbonate cages and fed with sterilized food and autoclaved water *ad-libitum*. At the end of experiment (6 wks), mice were euthanized by carbon dioxide asphyxiation and blood, bones and lymphoid tissues were harvested for further analysis. All the procedures were performed in accordance with the principles, recommendation and after the due approval of protocol submitted to Institutional Animal Ethics Committee of AIIMS, New Delhi, India (196/IAEC-1/2019).

### Scanning Electron Microscopy (SEM)

SEM was performed for scanning the surface of femur cortical region of bones. Briefly, for 2-3 days bone samples were stored in 1%-Triton-X-100 and later on bones were transferred in 1X PBS buffer till the analysis was carried out. After preparation of bone slices, samples were dried and sputter coating was performed. Subsequently, bones were scanned and imaged using Quanta 200 FEG SEM. SEM images were digitally photographed at 50,000 X magnification to capture the best cortical region of bones. The SEM images were further analysed through MATLAB software (Mathworks, Natick, MA, USA).

### Micro-Computed Tomography (μ-CT) and Bone Mineral Density (BMD) measurements

μ-CT scanning and BMD analysis were performed as described previously by Dar et al., 2018a, Dar et al., 2018b, Dar et al., 2018c). Briefly, after placing all the samples at correct orientation, scanning was carried out at 50 kV, 201 mA using 0.5 mm aluminium filter and exposure was set to 590 ms. For reconstruction of images NRecon software was used. For trabecular region analysis, ROI was drawn at a total of 100 slices in secondary spongiosa at 1.5mm from distal border of growth plates excluding the parts of cortical bone and primary spongiosa. For measuring and calculating the micro architectural parameters of bone samples CTAn software was employed. Several 3D-histomorphometric indices were obtained such as Bone volume/tissue volume ratio (BV/TV); trabecular thickness (Tb. Th); trabecular number (Tb. No.); Connectivity density (Conn. Den); trabecular separation (Tb. Sep.); Trabecular pattern factor (Tb. Pf.); total cross-sectional area (Tt. Ar.); Total cross-sectional perimeter (T. Pm); Cortical bone area (Ct. Ar); Bone perimeter (B. Pm); Average cortical thickness (Ct. Th). Volume of interest (VOI) of μ-CT scans was used to calculate the BMD of Lumbar vertebrae-5 (LV-5), Femoral and Tibial bones. BMD was measured by using hydroxyapatite phantom rods of 4 mm diameter with known BMD (0.25 g/cm3 and 0.75 g/cm3) as calibrator.

### Enzyme Linked Immunosorbent Assay (ELISA)

ELISA was carried out for quantitative estimation cytokines viz. IL-10, IL-17 and TNF-α in blood serum using commercially available kits as per the manufacturer’s instructions.

### Statistical Analysis

Statistical differences between sham and Ovx mice groups were assessed by using analysis of variance (ANOVA) with succeeding comparisons via student t-test paired or unpaired as appropriate. We performed analysis of significance in Sigma Plot (Systat Software, Inc., Germany). All the data values are articulated as Mean ± SEM (n=6). Statistical significance was determined as p ≤ 0.05 (*p < 0.05, **p < 0.01, ***p < 0.001) with respect to indicated group.

## Results

### Generation/Induction of IL-10 producing Bregs

In the present study, we for the first time have made an attempt to establish the role of regulatory B cells (Bregs) in modulating osteoclastogenesis. To determine whether Bregs possess the potential to modulate osteoclasts differentiation, we first induced Bregs differentiation under *in vitro* conditions. Bregs were thus generated as per established literature (Yanaba 2009; Wang et al., 2019). First, B cells were purified from splenocytes of mice via negative selection through magnetic beads (Fig.1A). After assessing the purity of isolated B cells (>95 %), we stimulated these purified B cells with lipopolysaccharide (LPS) (10ug/ml) for different time periods (5 and 24 hr). Since, Bregs have already been reported to function via secretion of their signature cytokine IL-10, thus we too determined the frequency of CD19^+^IL-10^+^ Bregs in our study. Interestingly, the frequency of total IL-10 producing B cells i.e., CD19^+^IL-10^+^ Bregs was significantly enhanced by LPS stimulation in a time dependent manner in comparison to control group (p < 0.001) (Fig. 1B). Of note, B10 Bregs with characteristic phenotype of CD19^+^CD1d^hi^CD5^+^IL-10^+^ were also significantly enhanced after LPS stimulation with respect to control group (Fig. 1C). These results of ours thus established successful generation of IL-10 producing Bregs in our culture conditions.

**Fig. 1.**
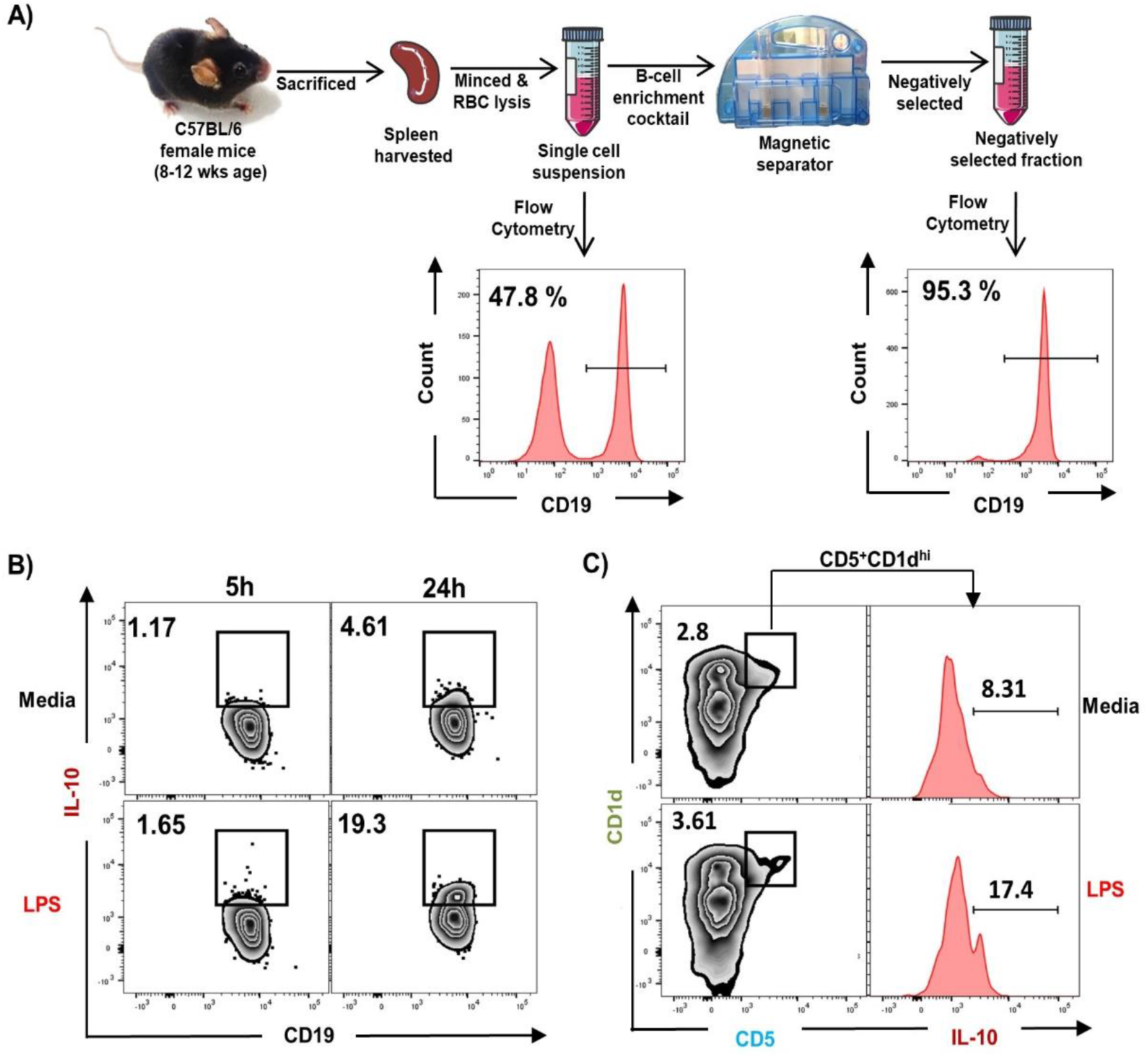
LPS activation promotes Bregs differentiation: **A)** Spleen was harvested and processed for negative selection of CD19^+^ B cells. After estimation of B cells purity (>95 %), cells were activated with LPS for different time periods. Assessed for B cells differentiation into Bregs, cells were analysed for Breg markers B) Zebra plot depicting the percentages of CD19^+^IL-10^+^ in media and LPS after 5 hour and 24 hours of stimulation C) Zebra plot depicting the percentages of CD19^+^CD5^+^CD1d^hi^ Bregs and histograms depicting IL-10 production by these cells. The results were evaluated by ANOVA with subsequent comparisons by Student t-test for paired or nonpaired data. Values are reported as mean ± SEM. Similar results were obtained in two different independent experiments with n=6. Statistical significance was considered as p≤0.05 (*p≤0.05, **p≤0.01, ***p≤0.001) with respect to indicated groups. (Mouse Image courtesy: Ms. Leena Sapra).

### Bregs inhibit differentiation of osteoclasts

Next, we wanted to investigate whether Breg cells exhibit the potential to suppress osteoclasts differentiation from bone marrow cells (BMCs) under *in vitro* conditions. To confirm this, we co-cultured BMCs with LPS induced Bregs (LPS-B) at different ratios (BMC: Bregs:: 10:1, 5:1 and 1:1) in the presence of M-CSF (30 ng/ml) and RANKL (100 ng/ml). After incubation for 4 days, cells were fixed and stained for TRAP to identify differentiated multinucleated osteoclasts (Fig.2A). Surprisingly, we observed that BMCs co-cultured with LPS activated Bregs at different cell ratios showed significant reduction (3-fold) in TRAP positive osteoclasts differentiation (Fig.2B-D). Moreover, area measurement analysis of TRAP positive multinucleated osteoclasts using Image J software too showed significant reduction (30 folds) in the area of osteoclasts in treated groups (Fig.2E). These results of ours thereby clearly suggest that Bregs inhibit RANKL induced osteoclastogenesis in a dose dependent manner. Further to dissect whether the suppressive effect of Bregs is mediated via either cell contact dependent (soluble factors) or cell contact-independent manner, we co-cultured BMCs and Bregs in trans-wells that prohibits direct cellular interactions. Remarkably, Bregs were found to significantly inhibit osteoclastogenesis in a dose dependent manner (5-fold) even in a trans-well setup thereby clearly establishing the role of soluble factors in mediating inhibitory activity of Bregs (Fig. 3A-E). Altogether, our data clearly establish that Breg inhibit differentiation of osteoclasts via mechanisms that may involve soluble molecules.

**Fig 2:**
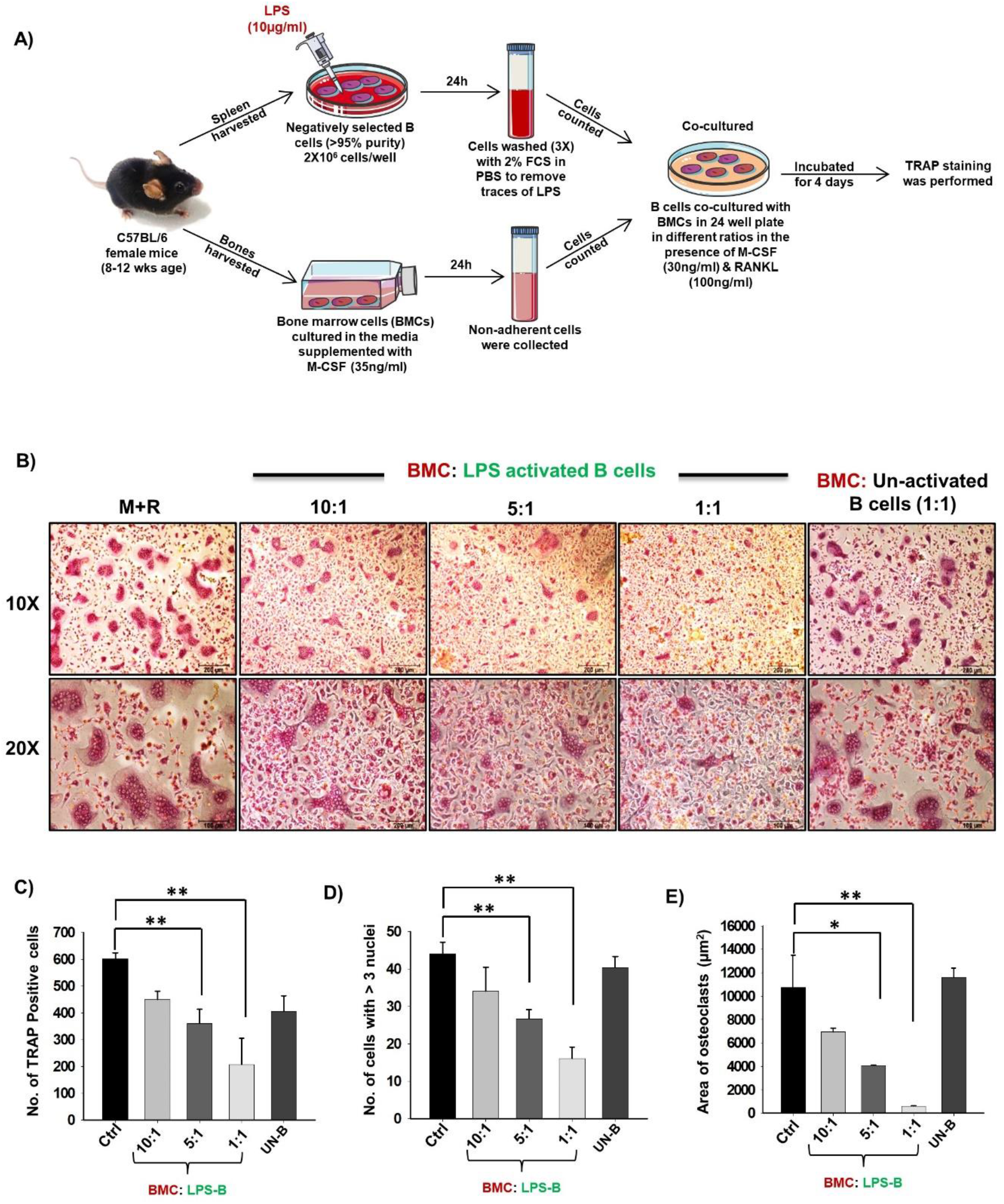
Inhibitory action of activated Breg cells on osteoclastogenesis (osteoclasts formation): **A)** BMCs and LPS activated and non-activated B cells were co-cultured in cell culture plate (cell-contact) in the presence of M-CSF (30 ng/ml) and RANKL (100 ng/ml) for 4 days. B cells were activated with LPS (10 ug/ml) for 24 hrs prior to co-cultures. B) LPS activated Bregs significantly inhibited the generation of multinucleated osteoclasts in a concentration dependent manner. C) Graphical representation depicting the number of TRAP positive cells. D) Bar graphs representing number of cells with more than three nuclei E) Bar graphs representing the area of multinucleated TRAP positive cells. Here, control denotes co-cultures of BMCs and B cells (non-activated). The results were evaluated by ANOVA with subsequent comparisons by Student t-test for paired or nonpaired data. Values are reported as mean ± SEM. Similar results were obtained in two different independent experiments with n=6. Statistical significance was considered as p≤0.05 (*p≤0.05, **p≤0.01, ***p≤0.001) with respect to indicated groups (Mouse Image courtesy: Ms. Leena Sapra).

**Fig 3:**
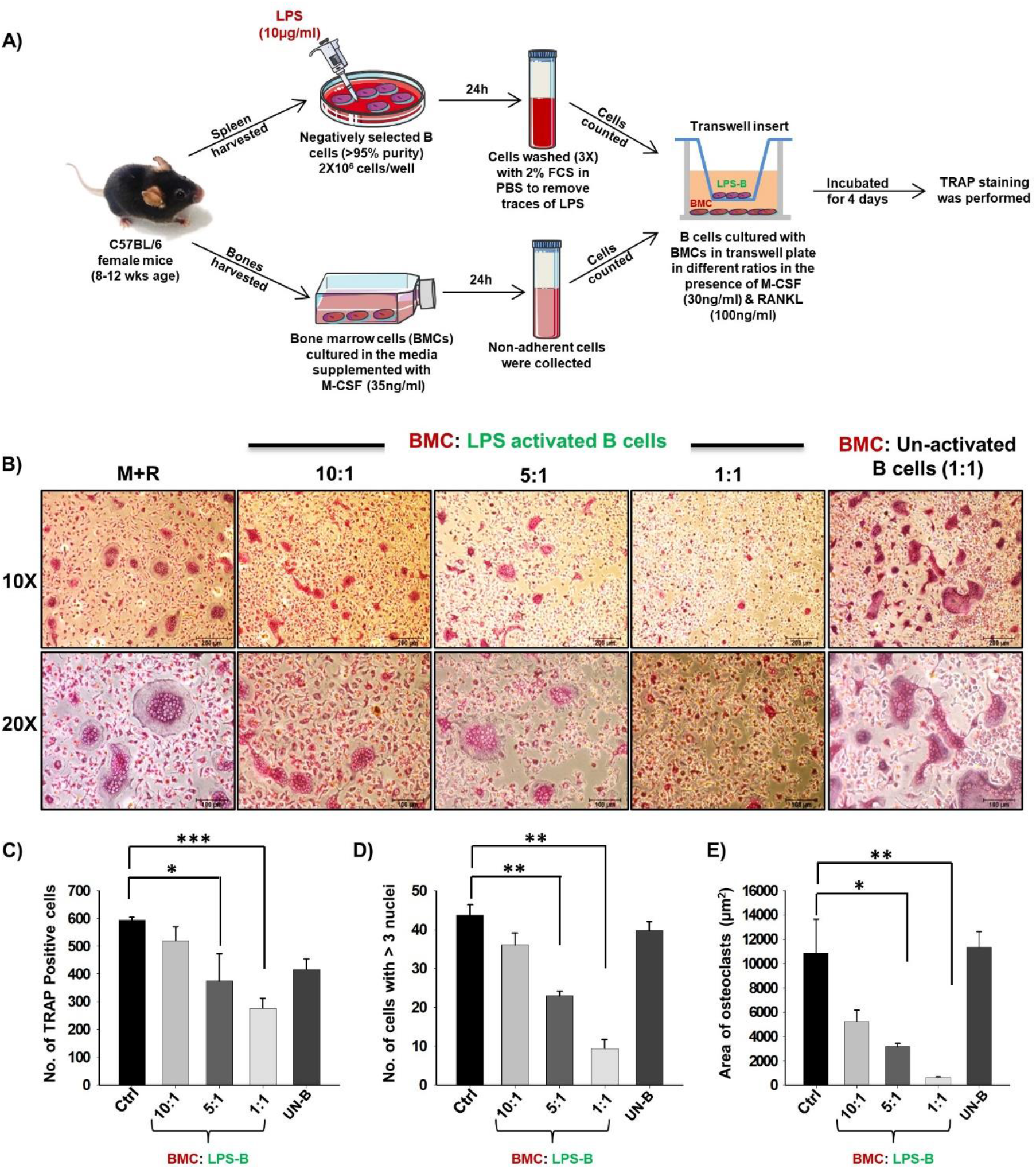
Inhibitory action of activated Breg cells on osteoclastogenesis (osteoclasts formation) in cell contact independent manner: **A)** BMCs and LPS activated and non-activated B cells were co-cultured in transwell insert (cell-contact) in the presence of M-CSF (30 ng/ml) and RANKL (100 ng/ml) for 4-5 days. B cells were activated with LPS (10 ug/ml) for 24 hrs prior to being added to co-cultures. B) LPS activated Bregs significantly inhibited the generation of multinucleated osteoclasts in a concentration dependent manner. C) Graphical representation depicting the number of TRAP positive cells. D) Bar graphs representing number of cells with more than three nuclei E) Bar graphs representing the area of multinucleated TRAP positive cells. Here, control denotes co-cultures of BMCs and B cells (non-activated). The results were evaluated by ANOVA with subsequent comparisons by Student t-test for paired or nonpaired data. Values are reported as mean ± SEM. Similar results were obtained in two different independent experiments with n=6. Statistical significance was considered as p≤0.05 (*p≤0.05, **p≤0.01, ***p≤0.001) with respect to indicated groups (Mouse Image courtesy: Ms. Leena Sapra).

### Bregs inhibits Osteoclast differentiation via IL-10 secretion

Various studies have shown that LPS-activated B cells expresses IL-10 cytokines, responsible for mediating its immunosuppressive activities (Yanaba et al., 2009, Kim et al., 2015). Also, among various subtypes of Bregs, CD19^+^CD1d^hi^CD5^+^ (B10) Bregs are known to play a crucial role in preventing immunopathogenesis associated with various autoimmune diseases. Thus, it may be possible that higher expression of IL-10 by these Bregs could contribute towards inhibition of osteoclastogenesis in our cultures. Therefore, to ensure the relevance of IL-10 in Bregs mediated suppression of osteoclastogenesis, we next assessed the percentages of these CD19^+^IL-10^+^ and CD19^+^CD1d^hi^CD5^+^IL-10^+^ B10 Bregs in our LPS induced Bregs population via flow cytometric analysis. Intriguingly, we observed that percentages of all of these respective IL-10 producing Breg populations was significantly enhanced in LPS induced group with respect to control group (Fig.4A-I). Thus, our results for the first time establish the direct role of Bregs in inhibiting osteoclasts differentiation.

**Fig 4:**
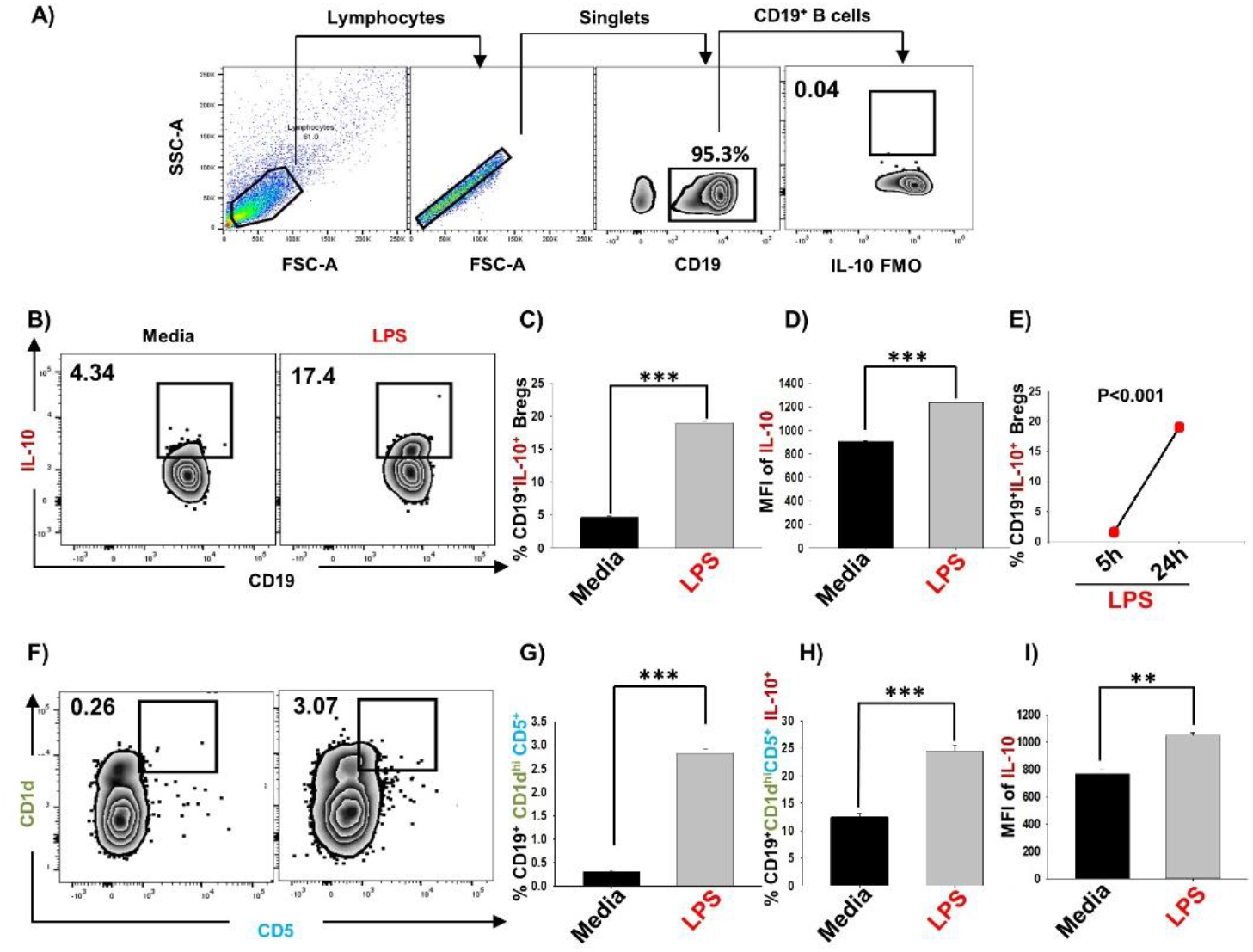
LPS activated B cells suppress osteoclastogenesis via expressing IL-10: After 24 hours of LPS stimulation cells were harvested and evaluated for estimating the expression of markers that may be associated with anti-osteoclastogenic function of B cells. A) Gating strategy followed for data analysis. B) Zebra plot depicting the percentages of CD19^+^IL-10^+^ Bregs in media and LPS activated B cells C) Graphical representations depicting percentages of CD19^+^IL-10^+^ Bregs D) MFI of IL-10 E) Line plot showing CD19^+^IL-10^+^ B cells after 5hr and 24 hr of stimulation. F) Zebra plot highlighting the percentages of CD19^+^CD1d^hi^CD5^+^ (B10) Bregs in media and LPS activated B cells. G) Graphical representation of CD19^+^CD1d^hi^CD5^+^ Breg H) Bar graph representing CD19^+^CD1d^hi^CD5^+^IL-10^+^Breg I) MFI of IL-10 on CD19+CD1dhiCD5+. The results were evaluated by ANOVA with subsequent comparisons by Student t-test for paired or nonpaired data. Values are reported as mean ± SEM. Similar results were obtained in two different independent experiments with n=6. Statistical significance was considered as p≤0.05 (*p≤0.05, **p≤0.01, ***p≤0.001) with respect to indicated Sham group.

### Successful development of postmenopausal osteoporotic mice model

Next to investigate the likely contribution of regulatory B cells in modulating bone health under normal and osteoporotic conditions, we firstly developed and authenticated postmenopausal osteoporotic mice model, a prime requirement for our study. For accomplishing the same, female C57BL/6 mice were divided in two groups viz. Sham and Ovx. At the end of experiment, blood serum was analysed for estradiol levels and bones were harvested for assessing the effect of estrogen deficiency on bone loss (Fig. 5A). Our results clearly indicated significant reduction in estradiol levels from 29 pg/ml (sham) to 9 pg/ml (3-fold) in Ovx group (p < 0.01) (Fig. 5B). Thus, after successful development of postmenopausal osteoporotic mice model we next investigated the effect of estrogen loss on bone loss and histomorphometric parameters. Scanning electron microscopy (SEM) and micro-computed tomography (μ-CT) analysis (a gold standard for determining bone health) was next performed. SEM analysis of cortical region (femoral bone) demonstrated enhanced number of lacunae or resorption pits in Ovx group with respect to control group (Fig. 5C). To further analyse the SEM 2D-images in more statistical manner, we performed MATLAB (matrix-laboratory) analysis to determine the correlation between bone mass and bone loss in both the groups. MATLAB analysis specifies degree of homogeneity where red colour indicates higher correlation values (high bone mass) and blue colour symbolizes lower correlation values (more bone loss). Outcomes of MATLAB analysis, clearly points that Ovx group has lesser correlation values or higher bone loss in Ovx group in comparison to sham group (Fig. 5D).

**Fig 5:**
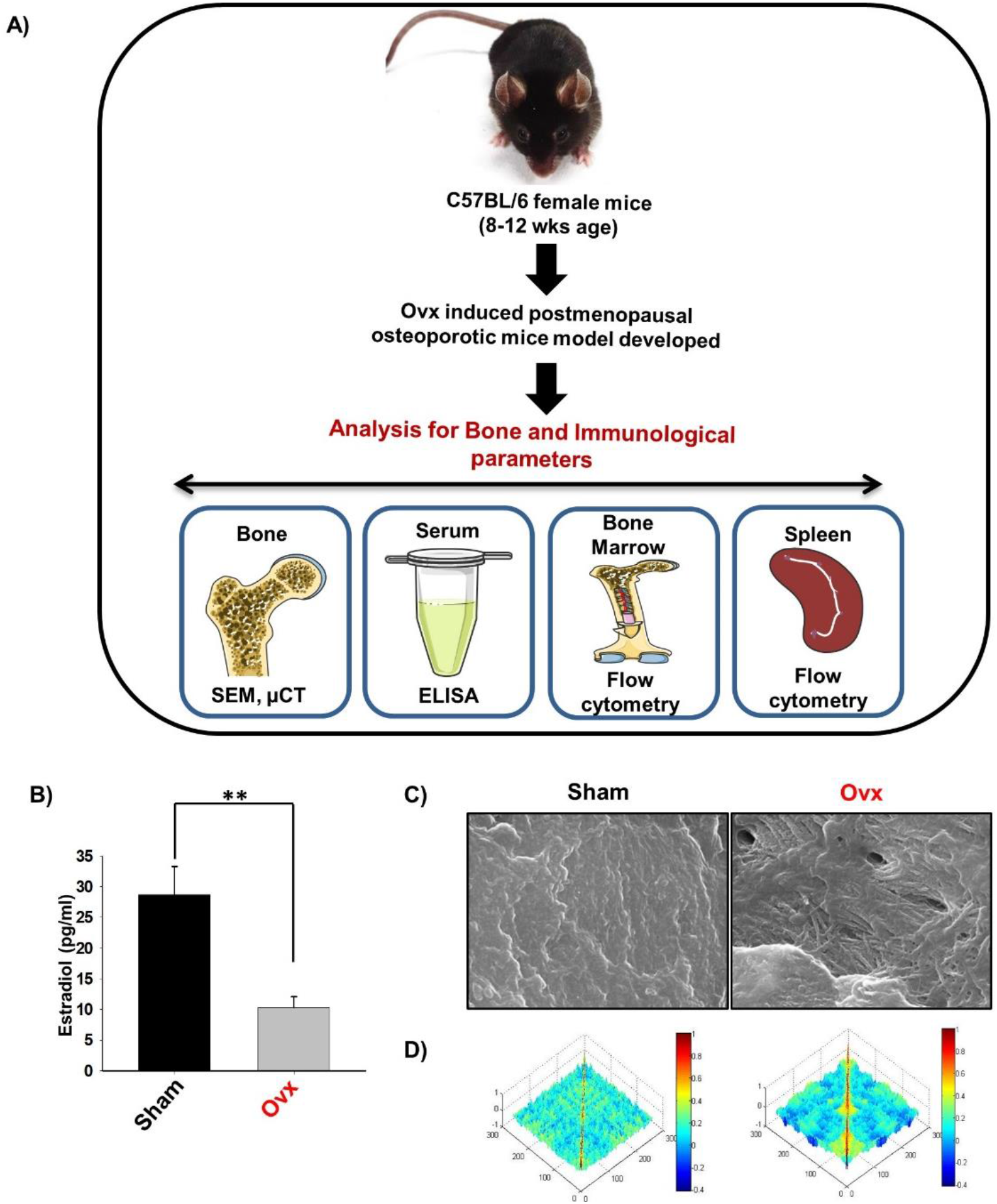
Estrogen deficiency augments bone loss in Ovx mice model. A) Experimental Layout B) Estrogen level was estimated in blood serum of sham and Ovx mice groups. Mice were sacrificed at the end of experiment and cortical bones of both the groups were collected for SEM analysis. C) 2D SEM images. D) 2D MATLAB analysis of SEM images. The above images are indicative of one independent experiment and comparable results were obtained in two different independent experiments with n=6 mice/group/experiment. The results were evaluated by ANOVA with subsequent comparisons by Student t-test for paired or non-paired data. Values are reported as mean ± SEM. Similar results were obtained in two different independent experiments with n=6. Statistical significance was considered as p≤0.05 (*p≤0.05, **p≤0.01, ***p≤0.001) with respect to indicated Sham group. (Mouse Image courtesy: Ms. Leena Sapra).

Moving ahead, we next authenticated the successful development of osteoporotic conditions in our model via μ-CT analysis. As Lumbar-vertebrae-5 (LV-5) is one of the peculiar regions to diagnose osteoporotic conditions (Sapra et al., 2021). Thus, we analysed the effect of estrogen deficiency on LV-5 trabecular region along with trabecular region of femoral and tibial bones. Interestingly, μ-CT data revealed that estrogen deficiency significantly impaired the micro-architecture of LV-5, femoral and tibial bones in Ovx group in comparison to sham group (Fig. 6A). Moreover, ameliorated bone loss condition in Ovx group is further supported by significantly reduced bone mineral density (BMD) in Ovx group with respect to sham group (Fig.6B). The quantitative analysis of trabecular and cortical bones which provides an overview of the utmost fundamental histomorphometric indices that were derived from bone micro-architecture 3D images is shown in Table 1. Altogether, our results clearly established the successful development of postmenopausal osteoporotic mice model.

**Fig 6:**
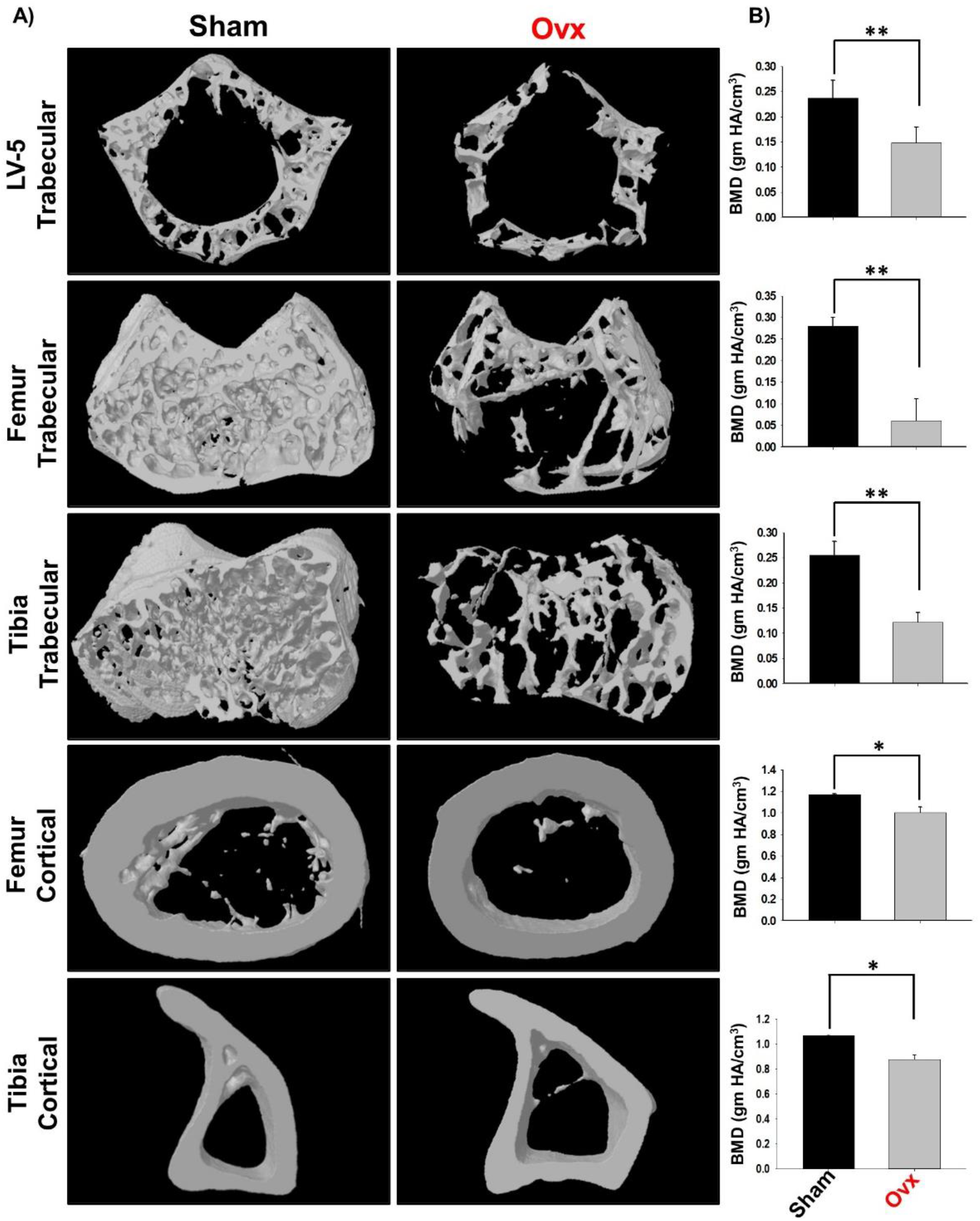
Estrogen deficiency deteriorates microarchitecture of trabecular and cortical regions of Lumbar Vertebrae-5 (LV-5), Femoral and Tibial bones. A) 3D micro-CT reconstructions of LV-5 trabecular region, Femur trabecular region, Tibia trabecular, Femur cortical and Tibia cortical regions of sham and Ovx groups B) Graphical representations of Bone Mineral Density (BMD) of LV-5 trabecular region, Femur trabecular region, Tibia trabecular, Femur cortical and Tibia cortical regions of sham and Ovx groups. The results were evaluated by ANOVA with subsequent comparisons by Student t-test for paired or non-paired data. Values are reported as mean ± SEM. Similar results were obtained in two different independent experiments with n=6. Statistical significance was considered as p≤0.05 (*p≤0.05, **p≤0.01, ***p≤0.001) with respect to indicated Sham group.

**Table 1:** Estrogen deficiency effects bone Histomorphometric parameters. Histomorphometric indices of LV-5 trabecular, Femur trabecular, Tibia trabecular, Femur cortical and tibia cortical of Sham and Ovx mice groups. Bone volume/tissue volume ratio (BV/TV); trabecular thickness (Tb. Th); trabecular number (Tb. No.); Connectivity density (Conn. Den); trabecular separation (Tb. Sep.); Trabecular pattern factor (Tb. Pf.); total cross-sectional area (Tt. Ar.); Total cross-sectional perimeter (T. Pm); Cortical bone area (Ct. Ar); Bone perimeter (B. Pm); Average cortical thickness (Ct. Th). The results were evaluated by ANOVA with subsequent comparisons by Student t-test for paired or nonpaired data. Values are reported as mean ± SEM. Similar results were obtained in two different independent experiments with n=6. Statistical significance was considered as p≤0.05 (*p≤0.05, **p≤0.01, ***p≤0.001) with respect to indicated Sham group.

| Bone Parameters | Sham | Ovx |
| --- | --- | --- |
| **LV5** |  |  |
| BV/TV (%) | 92.07 ± 2.84 | 60.87 ± 6.62** |
| Tb. Th (mm) | 0.06 ± 0.001 | 0.04 ± 0.001** |
| Tb. No (mm<sup>-1</sup>) | 11.8 ± 0.86 | 2.49 ± 0.08** |
| Conn. Den (mm<sup>-3</sup>) | 183.05 ± 43 | 89.66 ± 6.1 |
| Tb. Sp. (mm) | 0.01 ± 0.00 | 0.27 ± 0.003*** |
| Tb. Pf (mm<sup>-1</sup>) | 18.4 ± 0.22 | 29.7 ± 0.64* |
| **Femur Trabecular** |  |  |
| BV/TV (%) | 60.93 ± 3.21 | 8.24 ± 1.18** |
| Tb. Th (mm) | 0.068 ± 0.005 | 0.04 ± 0.003* |
| Tb. No (mm<sup>-1</sup>) | 13.6 ± 0.8 | 1.44 ± 0.18** |
| Conn. Den (mm<sup>-3</sup>) | 227 ± 29.6 | 58.4 ± 6.02** |
| Tb. Sp. (mm) | 0.07 ± 0.01 | 0.66 ± 0.03** |
| Tb. Pf (mm<sup>-1</sup>) | 11.12 ± 1.54 | 26.11 ± 0.80** |
| **Tibia Trabecular** |  |  |
| BV/TV (%) | 88.93 ± 0.09 | 5.23 ± 0.15*** |
| Tb. Th (mm) | 0.07 ± 0.001 | 0.05 ± 0.0006** |
| Tb. No (mm<sup>-1</sup>) | 13.1 ± 0.45 | 1.1 ± 0.15*** |
| Conn. Den (mm<sup>-3</sup>) | 60.24 ± 3.88 | 24.79 ± 0.95** |
| Tb. Sp. (mm) | 0.04 ± 0.002 | 0.53 ± 0.01*** |
| Tb. Pf (mm<sup>-1</sup>) | 15.14 ± 1.01 | 26.43 ± 0.78** |
| **Femur Cortical** |  |  |
| Tt. Ar (mm<sup>2</sup>) | 2.22 ± 0.05 | 1.5 ± 0.19* |
| T. Pm (mm) | 5.69 ± 0.09 | 4.95 ± 0.06* |
| Ct. Ar (mm<sup>2</sup>) | 0.85 ± 0.007 | 0.63 ± 0.04* |
| B. Pm (mm) | 10.33 ± 0.17 | 8.95 ± 0.15* |
| Ct. Th (mm) | 0.17 ± 0.001 | 0.12 ± 0.01* |
| **Tibia Cortical** |  |  |
| Tt. Ar (mm<sup>2</sup>) | 1.46 ± 0.012 | 1.23 ± 0.02** |
| T. Pm (mm) | 7.4 ± 0.79 | 5.7 ± 0.25 |
| Ct. Ar (mm<sup>2</sup>) | 0.70 ± 0.012 | 0.58 ± 0.02* |
| B. Pm (mm) | 10.4 ± 0.2 | 9.06 ± 0.25* |
| Ct. Th (mm) | 0.15 ± 0.001 | 0.11 ± 0.004** |

### IL-10 producing Bregs inhibit bone loss in osteoporotic mice model

Estrogen deficiency induced bone loss is mediated by various factors viz. immunological, biochemical etc. Also, the role of cytokine imbalance in regulating bone health is well established. Results from our lab (Dar et al., 2018a, Dar et al., 2018b) along with others have reported significantly reduced levels of osteoprotective cytokine IL-10 and higher levels of osteoclastogenic cytokines IL-17 and TNF-α under Ovx conditions (Fig. 7A-B). Extensive literature suggests that the major source of IL-10 is adaptive immune cells viz. Tregs and Bregs. Recently, several studies have reported that apart from Tregs, Bregs are also a major source of IL-10 in physiological conditions. Therefore, a strong possibility exists that reduction of IL-10 levels in osteoporotic conditions may not be primarily due to dysregulation of Tregs alone. Based on these studies we strongly believe and propose that alteration in Bregs population could also be one of the major contributing factors for observed significant reduction in IL-10 levels observed Ovx conditions. Since, Bregs are predominant source of IL-10 in various physiological conditions, we next asked whether the observed significantly reduced levels of IL-10 are due to dysregulation in Bregs population in Ovx mice? Since B10 (CD19^+^CD1d^hi^CD5^+^IL-10^+^) Bregs are the most established and studied subtype of Bregs, thus we made an attempt to elucidate the role of Bregs in osteoporotic conditions. We thus analysed the frequency of Bregs in both BM (prime site of osteoclastogenesis) and spleen (main site of Bregs induction) in mice. Interestingly, the frequencies of total IL-10 producing B cells i.e., CD19^+^IL-10^+^ cells were significantly reduced in Ovx group in comparison to control group (p < 0.01) in both BM and spleen (Fig. 8A-C). Notably, the population of CD19^+^CD1d^hi^CD5^+^IL-10^+^ B10 Bregs was also observed to be significantly reduced in both BM and spleen of Ovx group in comparison to sham group (p < 0.05) (Fig. 8D-H). Thus, our results provide novel insight in defining the phenotype and osteo-protective abilities of regulatory B cells in post-menopausal osteoporotic condition as both numerical and functional defects in Bregs could be responsible for development of inflammatory bone loss condition ion osteoporotic patients. Altogether, these results corroborate our in vitro findings thereby establishing the unexplored role of Bregs in osteoporosis.

**Fig 7:**
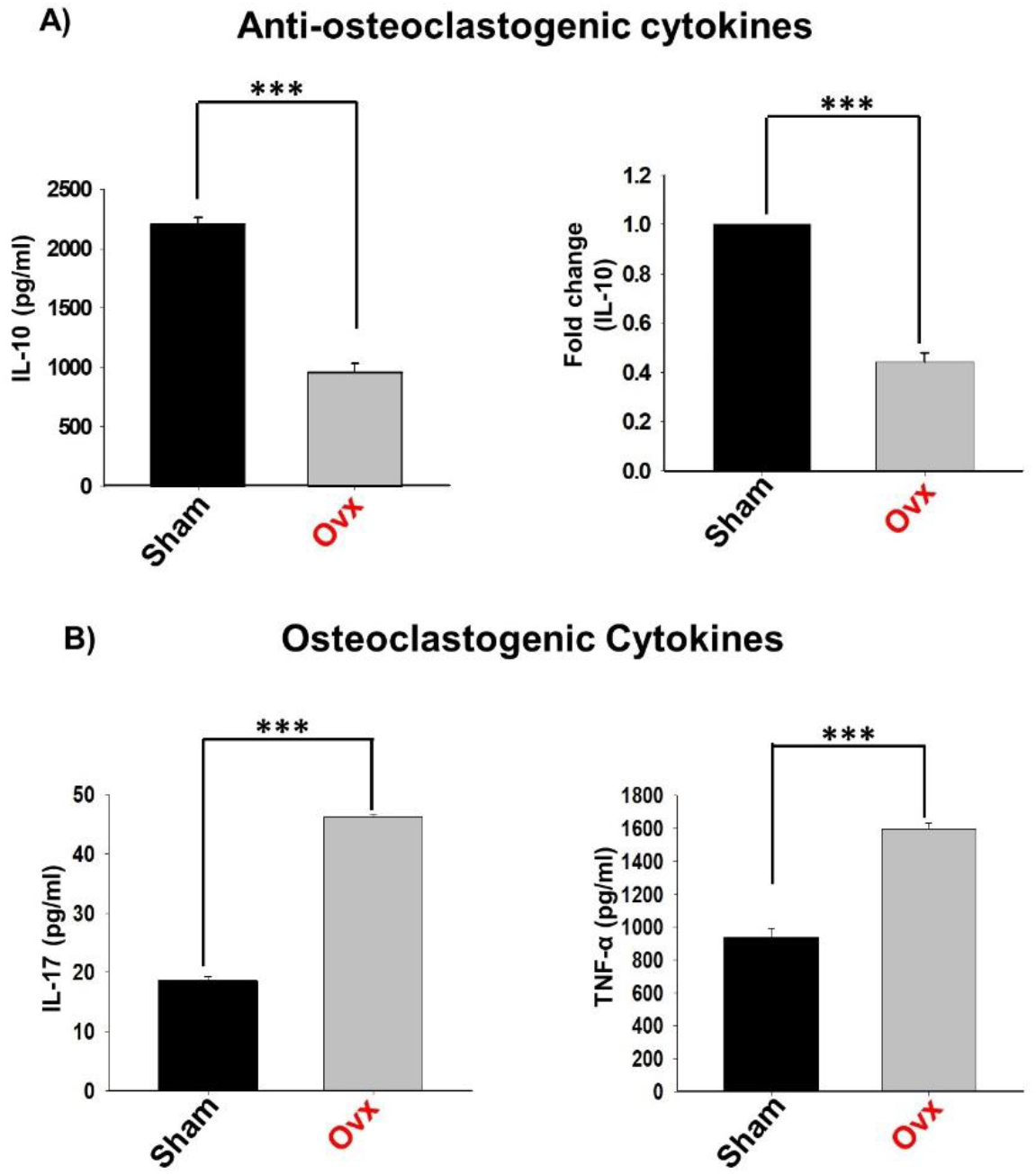
Deficiency of Estrogen skews osteoclastogenic and anti-osteoclastogenic cytokine levels in Ovx mice. Cytokines were analysed in serum samples of mice by ELISA. A) Levels of IL-10 cytokine B) Level of IL-17 in sham and Ovx groups. C) Level of TNF-α in sham and Ovx groups. The results were evaluated by using ANOVA with subsequent comparisons by Student t-test for paired or non-paired data, as appropriate. Values are expressed as mean ± SEM (n=6) and similar results were obtained in two independent experiments. Statistical significance was defined as p≤0.05, *p≤0.05, **p < 0.01 ***p≤0.001 with respect to indicated mice group.

**Fig 8:**
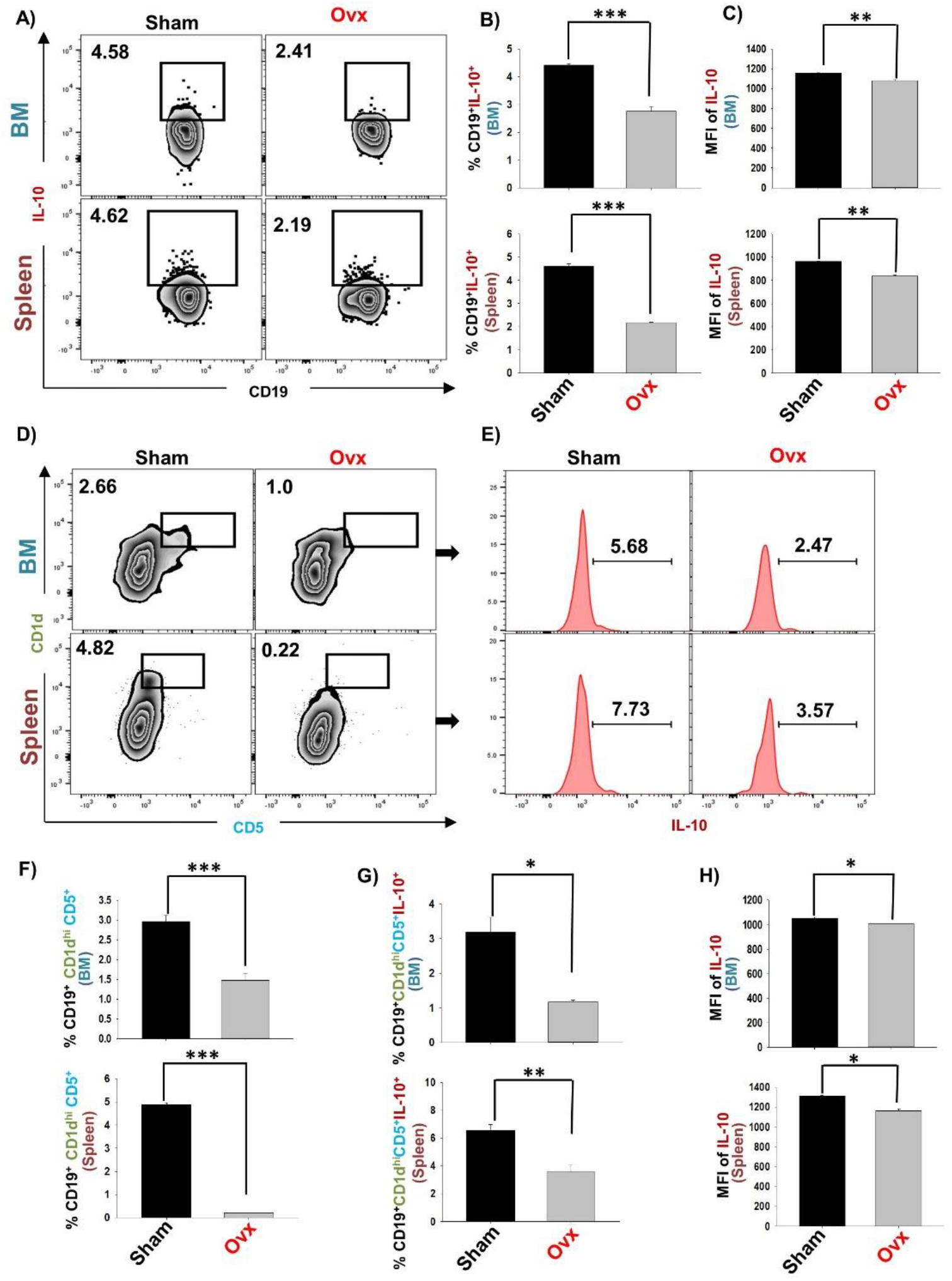
Reduction in Bregs population enhances the bone loss in Ovx mice: Cells from Bone Marrow (BM) and spleen of Sham and Ovx groups were harvested and analysed by flow cytometry for percentage of various Bregs subsets A) Zebra plot depicting percentages of CD19^+^IL-10^+^ B cells in BM and spleen of sham and Ovx B) Bar graph representing percentages CD19^+^IL-10^+^ B cells in sham and Ovx C) Bar graphs representing the MFI of IL-10. D) Zebra plot depicting percentages of CD19^+^CD1d^hi^CD5^+^ Bregs in BM and spleen E) Graphical representation of CD19^+^CD1d^hi^CD5^+^ B cells in sham and Ovx. F) Graphical representation of CD19^+^CD1d^hi^CD5^+^IL-10^+^ B cells in sham and Ovx. The results were evaluated by using ANOVA with subsequent comparisons by Student t-test for paired or non-paired data, as appropriate. Values are expressed as mean ± SEM (n=6) and similar results were obtained in two independent experiments. Statistical significance was defined as p≤0.05, *p≤0.05, **p < 0.01 ***p≤0.001 with respect to indicated mice group.

## Discussion

Osteoporosis is an inflammatory bone disease characterized by loss of estrogen at menopause. Estrogen loss during menopause leads to dysregulated bone remodelling process (Cline-Smith et al., 2020). Studies from our lab along with others have highlighted the relevance of immune system in maintaining bone health under physiological and osteoporotic conditions (Okamoto et al., 2017; Srivastava et al., 2018, Dar et al., 2018a; Yu et al., 2020). These studies laid foundation to the importance of immune system in osteoporosis i.e., Immunoporosis (Srivastava et al., 2018). Various studies demonstrated that the delicate balance between various inflammatory and anti-inflammatory cells is of vital importance for the maintenance of host integrity and skeleton system. Especially, the homeostatic balance between anti-inflammatory (Tregs) and inflammatory (Th17) cells is of utmost importance in various disease conditions including osteoporosis (Dar et al., 2018a, Shokeen et al., 2020; Sapra et al., 2021). Apart from Tregs there are also other regulatory lymphocytes with established immunosuppressive functions. Most important among these are the recently discovered Bregs.

The aim of the present study was to investigate the plausible role of Bregs in modulating bone loss via its effect on osteoclastogenesis. B cells that exhibit immunoregulatory potential have been observed in several disease models of autoimmunity such as RA, SLE, experimental autoimmune hepatitis (EAH) etc (Blair et al., 2010; Flores Borja et al., 2013; Liu et al., 2015). These Breg population via producing and secreting soluble cytokines (IL-10) with regulatory potential, or via expressing membrane bound surface molecules (TGF-β) promotes their regulatory functions. Nevertheless, mostly the regulatory mechanisms of Bregs are primarily based on the immunoregulatory cytokine IL-10. In 2008, Yanaba et al. identified a population of splenic CD19^+^CD1d^hi^CD5^+^ B cells that is highly enriched for IL-10 cytokine and named as B10 Bregs (Yanaba et al., 2008). Multiple studies demonstrated that IL-10 is an anti-osteoclastogenic cytokine that maintains bone health by inhibiting osteoclastogenesis (Evans et al., 2007; Park-Min et al., 2009). Moreover, significance of IL-10 is further highlighted in IL-10 deficient mice in which decreased bone mass, enhanced mechanical fragility and reduced bone formation was observed which are known to be the hallmarks of osteoporosis (Dresner et al., 2004). Moreover, Tregs via secreting IL-10 cytokine and along with other cytokines viz. IL-4 and membrane bound receptors such as cytotoxic T lymphocyte antigen (CTLA)-4 abrogates osteoclastogenesis (Zaiss et al., 2007). Although IL-10 is a pivotal immunomodulator in infectious diseases, autoimmune disorders, inflammatory bone loss conditions including osteoporosis, but no study till date have ever investigated the significance of IL-10 secreting Bregs in modulating osteoclastogenesis under *in vitro* and in osteoporotic mice model.

Thus, to understand the probable contribution of Bregs in modulating osteoclasts generation we carried out the present study. IL-10 is an established regulatory molecule of Bregs; but, expression of IL-10 cytokine is quite low in resting B cells thus to enhance the expression of IL-10, *in vitro* stimulation of B cells is required. Next, we activated purified B cells (>95 %) with LPS for different time periods (5 and 24 hr) and in consistent to previously reported studies we too observed a time dependent induction of IL-10 producing Bregs. Moving ahead, we further investigated whether, Breg cells can affect the formation of osteoclasts in vitro, we thus co-cultured LPS induced Bregs with bone marrow cells (BMCs) in different ratios for 4 days. Excitingly, we observed that Bregs inhibit the differentiation of osteoclasts from bone marrow cells in a dose dependent manner. We were further keen to know whether the observed suppression of osteoclastogenesis by Bregs in co-culture system is cytokine dependent (soluble factors) or require cell to cell contact for its anti-osteoclastogenic potential. Thus, we co-cultured LPS-Bregs cells and BMCs in transwell system for 4 days and our results clearly indicated that anti-osteoclastogenic potential of Bregs is primarily mediated by IL-10. Moreover, our flow cytometric data too confirmed that Bregs have enhanced induction of IL-10 which thereby inhibits osteoclastogenesis. Altogether, these findings of ours establish the role of Bregs in modulating bone health via suppression of osteoclastogenesis.

A study reported in 2013, by Flores et al. group demonstrated that numerical defect or profound reduction in frequencies of Bregs was found to be the causative factor for RA in humans (Flores Borja et al., 2013). But the role of Bregs in case of postmenopausal osteoporosis has never been reported. Thus, to further confirm and extend these findings under *in vivo* conditions we employed post-menopausal osteoporotic mice model. Our *in vivo* Estradiol levels, SEM and μ-CT data confirmed the successful development of osteoporotic mice model. Furthermore, our *in vivo* flow cytometric data clearly indicated towards a numerical defect in CD19^+^IL-10^+^ and CD19^+^CD1d^hi^CD5^+^IL-10^+^ Bregs in both BM (Prime site of osteoclastogenesis) and spleen (main site of Bregs). Also, our serum cytokine data, corroborates with our flow cytometric data with significant reduction of IL-10 levels in osteoporotic mice. Taken together, our data unravel and establish the unexplored role of Bregs in case of osteoporosis (Fig.9). Our present study thus for the first time establish that Bregs exhibits anti-osteoclastogenic potential and reduction in Bregs number may contribute towards inflammatory bone loss observed in postmenopausal osteoporotic mice model. Thus, Bregs appear to be as potential candidates which can be targeted and employed as a cellular therapeutic tool for treating inflammatory bone loss observed in osteoporosis and provide a novel insight into Bregs biology in the context of postmenopausal osteoporosis.

**Fig 9:**
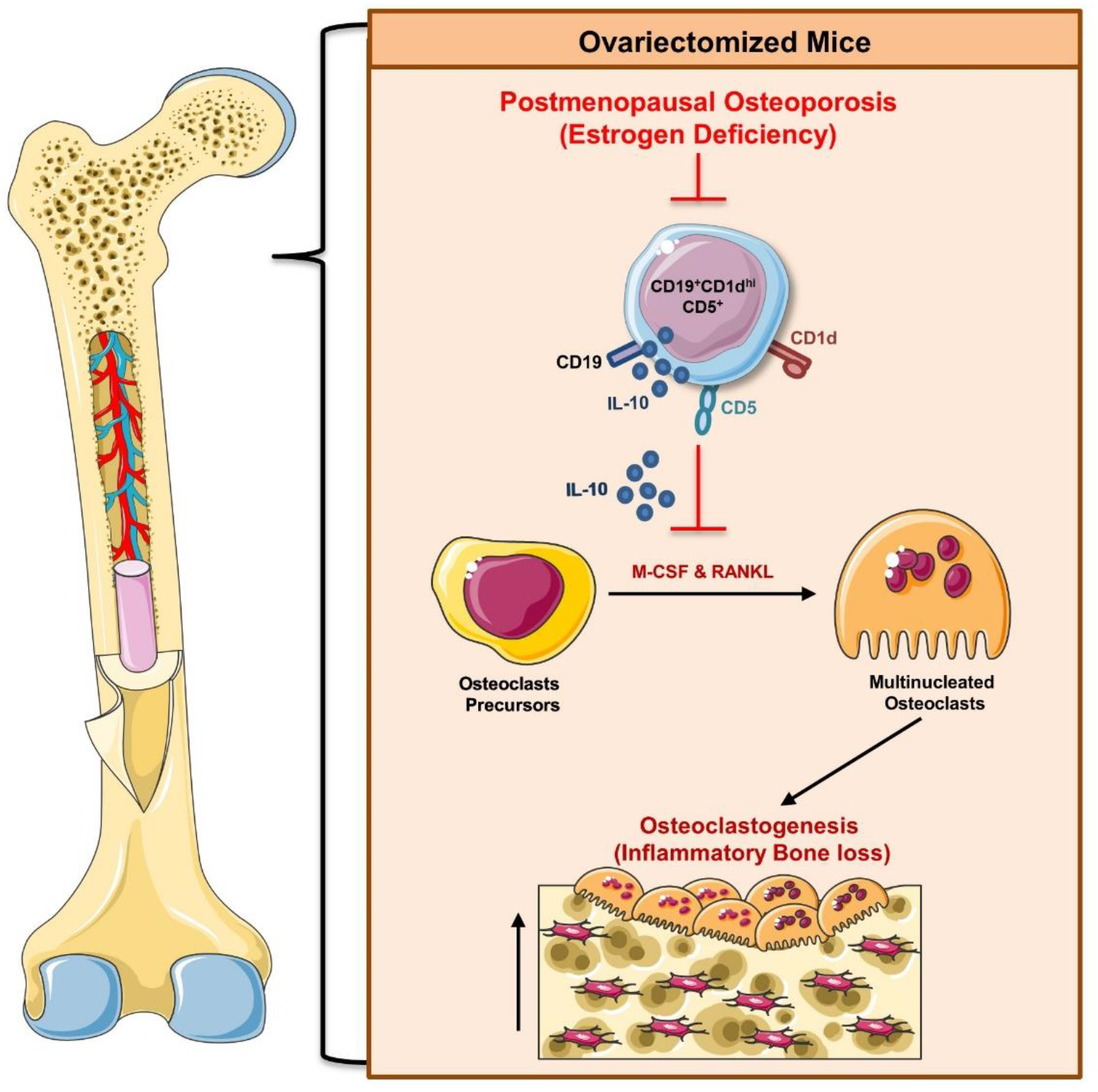
Summary of our results: LPS activated regulatory B cells exhibits the potential of inhibiting osteoclasts formation under *in vitro* conditions via expressing *in vitro*. Under *in vivo* conditions, deficiency of estrogen hormone leads to reduction in various subsets of Bregs: CD19^+^ CD1d^hi^CD5^+^ Bregs that further causes reduction in IL-10 cytokine levels thus leads to enhancement in osteoclastogenesis in postmenopausal osteoporotic mice model. (Image illustrated using Medical Art https://smart.servier.com/).

## Abbreviations

Bregs: Regulatory B cells
RANKL: Receptor activator of nuclear factor kappa B
TLR: Toll like receptor
LPS: Lipopolysaccharide
IL: Interleukin
TGF: transforming growth factor
RA: Rheumatoid arthritis
SLE: Systemic lupus erythematosus
EAE: Experimental autoimmune encephalitis
PMA: Phorbal 12-myristate 13-acetate
BMCs: Bone marrow cells
M-CSF: Monocyte colony stimulating factor
TRAP: Tartrate acid phosphatase
BMD: Bone mineral density
SEM: Scanning electron microscopy
Tregs: Regulatory T cells
CIA: Collagen induced arthritis

## Acknowledgment

This work was financially supported by projects: DST-SERB (EMR/2016/007158), Govt. of India and intramural project from All India Institute of Medical Sciences (AIIMS, A-798), New Delhi-India sanctioned to RKS. LS, PKM, BV and RKS acknowledge the Department of Biotechnology AIIMS, New Delhi-India for providing infrastructural facilities. The authors thank CCRF shared FACS Facility, AIIMS for acquisition of samples. LS thank UGC for research fellowship and AB thank DST SERB project for research fellowship.

## Author contributions

RKS contributed to conceptualization and investigation of study. LS and AB contributed to methodology and formal analysis of data. PKM carried out cytokine analysis. RKS and LS contributed in writing and editing of manuscript. BV provided valuable inputs in the study design. All authors reviewed the manuscript.

## Compliance with ethical standards

All applicable institutional and/or national guidelines for the care and use of animals were followed.

